# Loss of FBXO9 enhances proteasome activity and promotes aggressiveness in acute myeloid leukemia

**DOI:** 10.1101/732198

**Authors:** R. Willow Hynes-Smith, Samantha A. Swenson, Heather Vahle, Karli J. Wittorf, Mika Caplan, Catalina Amador, R. Katherine Hyde, Shannon M. Buckley

**Author notes:** Address Correspondence To: Shannon M. Buckley, Ph.D., Department of Genetics, Cell Biology, and Anatomy, University of Nebraska Medical Center, 985805 Nebraska Medical Center, Omaha, NE 68198.

## Abstract

The hematopoietic system is maintained throughout life by hematopoietic stem cells that are capable of differentiating into all hematopoietic lineages. An intimate balance between self-renewal, differentiation, and quiescence is required to maintain hematopoiesis. Disruption of this balance can result in hematopoietic malignancy, including acute myeloid leukemia (AML). FBXO9, from the F-box E3 ubiquitin ligases, is down-regulated in patients with AML compared to normal bone marrow. FBXO9 is the substrate recognition component of the Skp1-Cullin-F-box (SCF)-type E3 ligase complex. *FBXO9* is highly expressed in hematopoietic stem and progenitor populations, which contain the tumor-initiating population in AML. In AML patients, decrease in *FBXO9* expression is most pronounced in patients with the inversion of chromosome 16 (inv(16)), a rearrangement that generates the transcription factor fusion gene, *CBFB-MYH11*. To study FBXO9 in malignant hematopoiesis, we generated a conditional knockout mouse model using a novel CRISPR/Cas9 strategy. Our data show that deletion of *Fbxo9* in mice expressing *Cbfb-MYH11* leads to markedly accelerated and aggressive leukemia development. In addition, we find loss of FBXO9 leads to increased proteasome expression and tumors are more sensitive to bortezomib suggesting that FBXO9 expression may predict patient response to bortezomib treatment.

## INTRODUCTION

Acute myeloid leukemia (AML) is a hematologic malignancy resulting in an accumulation of immature myeloid blasts impairing normal hematopoiesis^1^. This disease accounts for 35% of new leukemia diagnoses and 48% of leukemia-related deaths^2^. In 2019, AML is estimated to be the most frequently diagnosed leukemia and the only common form with a higher mortality rate than incidence. Although new therapies have recently been approved, they are limited to specific subtypes of AML and only increase the treatment options for limited subsets of patients^3–6^. To develop more effective and less toxic therapies for use across multiple subtypes, we must better understand the mechanisms underlying development and progression of AML.

One system that has not been extensively studied in the context of AML is the ubiquitin proteasome system (UPS). The UPS coordinates the degradation of proteins globally and compartmentally within a cell and is a key regulatory mechanism for many cellular processes including cell cycle, transcription, and proliferation^7^. Two main components of this system are the E3 ubiquitin ligases that determine substrate specificity and the 26S proteasome responsible for protein degradation. This system has proven to be a viable target in hematologic malignances either through targeting the 26S proteasome with drugs such as bortezomib^8^ or by altering substrate recognition of E3 ligases as is done with thalidomide^9^. The successful utilization of these drugs in other hematologic malignancies suggests that targeting the UPS in AML could prove effective in treating the disease.

Ubiquitin E3 ligases can be classified as RING-finger, HECT-domain, or RBR based on their domains and mode of transferring ubiquitin to the substrate^10^. The largest family of E3 ligases, the Skp1-Cul1-F-box (SCF) family, is composed of a core complex including S-phase kinase-associated protein 1 (Skp1) and Cullin 1 (Cul1) which are scaffolding proteins that bring the ubiquitin binding RING-finger protein Rbx in proximity with the substrate recognition F-box protein component^11^. There are 69 F-box proteins that interact with Skp1 via the F-box domain and with substrate proteins through a variety of substrate-recognition domains^12,13^. Dysregulation of E3 ligases, including many F-box proteins, has been correlated with aberrant hematopoiesis and malignant transformation^14,15^. Many proteins from the F-box family have been classified as tumor suppressors, others as oncogenes, and a third set with context-specific roles^16^. FBXW7, for example, plays an integral role in hematopoietic stem and progenitor cell (HSPC) self-renewal and its loss has been linked to drug resistant T-cell acute lymphoblastic leukemia, whereas loss in chronic myeloid leukemia (CML) inhibits initiation and progression of disease^17–19^. Loss of FBXO4 increases extramedullary myeloid hematopoiesis and is highly expressed in various lymphomas^20^. Other family members have been linked to leukemia cell proliferation, including FBXL2, FBXL10, FBXW11, and SKP2^21–24^. Furthermore, overexpressed FBXO9 in multiple myeloma (MM) degraded TEL2 and TTI1, shifting signaling from mTORC1 to mTORC2, thus causing increased proliferation and survival^25^.

In this study, we identified FBXO9 as an important regulator of AML and found it has low expression in patients across all AML subtypes. To study the role of *Fbxo9* in AML, we developed a conditional knockout (cKO) mouse model and monitored the leukemia response *in vivo*. We utilized a mass spectrometry (MS)-based approach to identify proteins upregulated when *Fbxo9* expression is lost in tumors and identified various proteins previously shown to participate in cancer-related mechanisms such as metastasis, proliferation, invasion, and metabolism. Of particular interest, we found that many upregulated proteins participate in proteasome-regulated pathways. Furthermore, our *in vitro* analysis found that cKO tumors had increased proteasome activity and responded better to bortezomib treatment. Our studies provide insight into the role of the UPS in AML and present evidence that selecting patients according to *FBXO9* expression could be used as a method of identifying tumors to treat with proteasome inhibitors.

## RESULTS

### FBXO9 has low expression in AML and expression correlates to poor survival

To identify F-box proteins involved in initiation and/or progression of AML, we analyzed patient data from the Leukemia MILE study for expression of 61 F-box proteins^26^. Analysis revealed that *FBXO9* has the lowest expression in inv(16), MLL-rearranged, and t(8;21) AMLs among the F-box proteins. Additionally, when compared to healthy bone marrow (HBM), CML and myelodysplastic syndrome, *FBXO9* showed decreased expression (Figure 1A, S1A). Further analysis across a wider variety of AML subtypes, including normal and complex karyotype and t(15;17), revealed that *FBXO9* is consistently down-regulated across all subtypes (Figure 1B). As AML is the second most common childhood leukemia, we utilized the TARGET pediatric study to analyze *FBXO9* expression and again found down-regulation of FBXO9 in all AML sub-types except patients with normal karyotype (Figure 1C)^27^.

**Figure 1.**
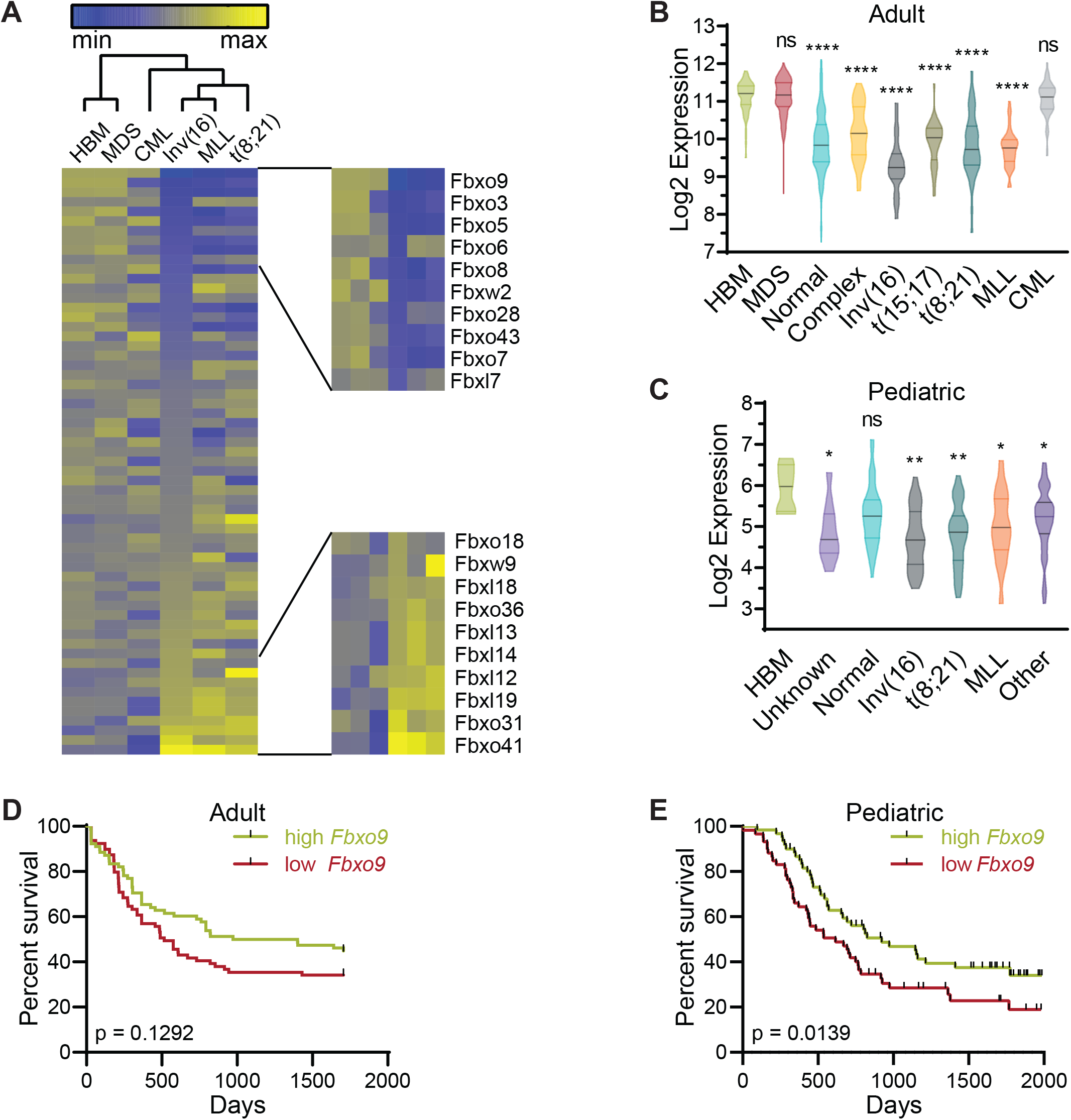
*Fbxo9* expression is reduced in AML patients and correlates with poor survival. A Of all F-box family proteins, *FBXO9* has the lowest expression in AML subtypes. B-C Analysis of patient samples from the B adult MILE and C pediatric TARGET studies reveals that AML patients have low *FBXO9* expression across a variety of subtypes when compared to healthy BM. D-E Poor survival correlates with low *FBXO9* expression in both D adults and E children (* p < 0.05, ** p < 0.01, *** p < 0.001, **** p < 0.0001).

Correlation of expression versus survival in adult and pediatric patients revealed that adult patients with *FBXO9* expression below the median tend to have a worse prognosis and shorter time of survival compared to patients with expression above the median, although not significant (Figure 1D, S1B). However, analysis of pediatric patients within the first 2000 days post diagnosis demonstrated that low *FBXO9* expression correlated with poor survival (Figure 1E). Patients who went in remission and survived over 2000 days post initial diagnosis had no significant difference in survival with any of the sub-types (Figure S1C-H)^28^. Taken together, these findings suggest that *FBXO9* expression is decreased in AML cells and that low expression correlates with poor survival at early time-points from initial diagnosis.

### Generation of conditional knockout of Fbxo9

To study the role of *Fbxo9* in hematopoiesis and AML, we generated a cKO mouse model. FBXO9 has two known domains, the F-box domain for binding the SCF complex and the protein-binding TPR domain^11^. Using the Easi-CRISPR method, we introduced LoxP sites flanking *Fbxo9* exon 4, which contains the majority of the TPR domain (Figure 2A)^29^. The targeted mouse was bred with an Mx1-cre mouse in which cre expression in the hematopoietic system is induced by the Mx1 promoter through administration of Polyinosinic:Polycytidilic acid (Poly(I:C))^30^.

**Figure 2.**
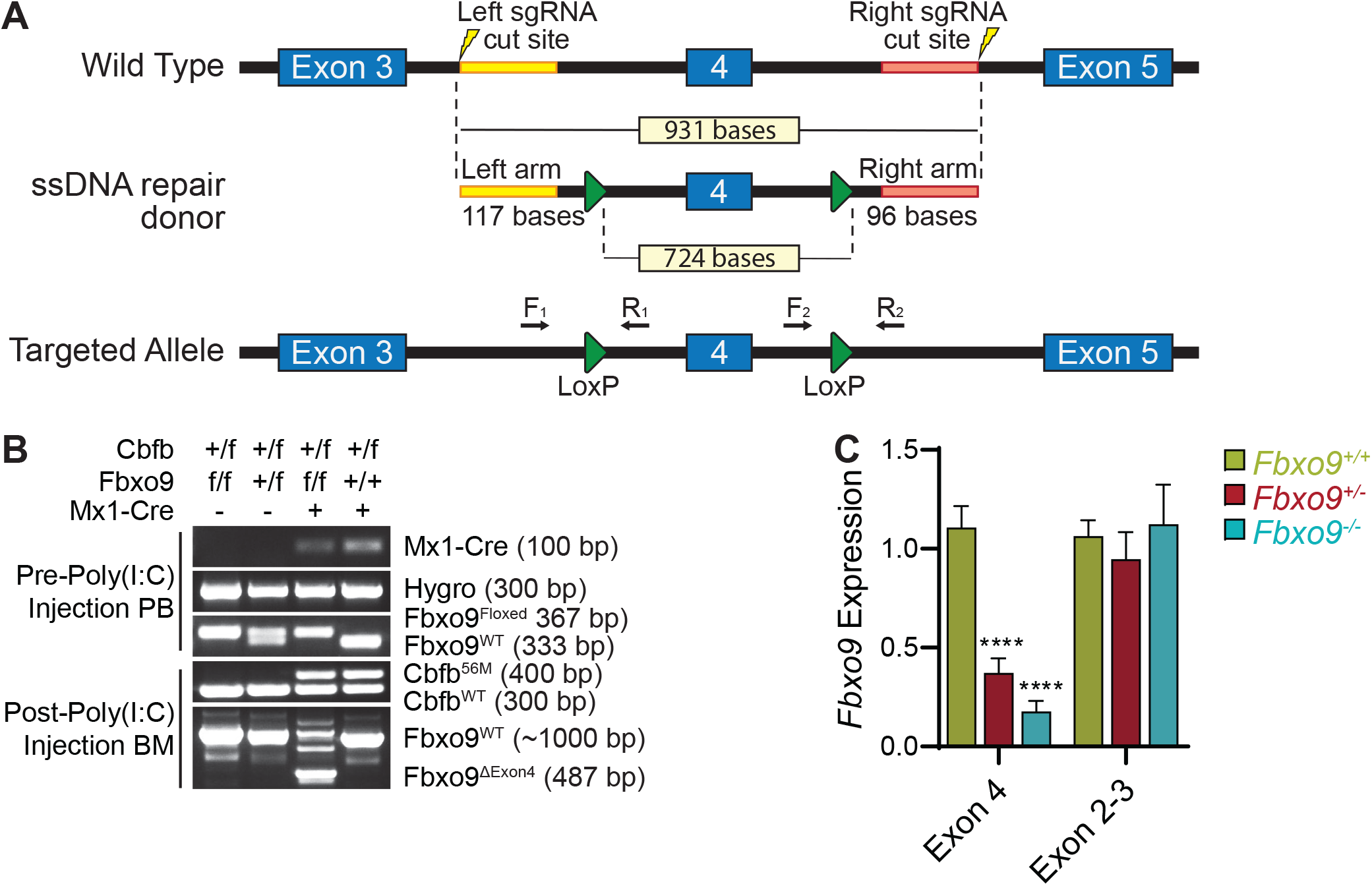
Generation of conditional *Fbxo9* knockout mouse model. A Using the Easi-CRISPR method, LoxP sites were introduced into the introns flanking *Fbxo9* exon 4. B Genotyping PCR demonstrates the introduction of LoxP sites and presence of *Cbfb^+/56M^* transgenic allele and shows the loss of exon 4 and expression of the *Cbfb-MYH11* fusion gene following injections with Poly(I:C). C *Fbxo9* exon 4 cKO confirmed by qRT-PCR analysis while exons 2-3 remain undisturbed (**** p < 0.0001).

We confirmed the genotypes by PCR and analyzed *Fbxo9* expression in replicate *Fbxo9^+/−^* and *Fbxo9^−/−^* mice compared to *Fbxo9^+/+^*. The expression within BM of our cKO groups was reduced ~55% *Fbxo9^+/−^* and ~80% *Fbxo9^−/−^* (Figure 2B-C). Loss of exon 4 results in ablation of TPR binding and initiates a frame-shift mutation leading to loss of the F-box domain and a premature stop as confirmed by PCR and sequencing in our mouse model (Figure S2A). Additionally, we transfected *Fbxo9* mutant plasmids into 293T cells. Mutants lacking the F-box or TPR domains showed a slight decrease in molecular weight whereas the mutant lacking bases corresponding to exon 4 showed a much greater decrease in molecular weight (Figure S2B). The band size correlated with the predicted molecular weight of a protein arising from cKO cells. This evidence indicates that deletion of exon 4 in our mouse model results in a truncated protein lacking the TPR and F-box domains.

### Fbxo9 deletion leads to alterations in HSPC populations

By analyzing different hematopoietic populations isolated from BM of a healthy WT mouse, we found that *Fbxo9* is most highly expressed in short- and long-term HSCs as well as some myeloid lineages (Figure 3A). To study the role of *Fbxo9* in normal hematopoiesis, we deleted *Fbxo9* by 3 sequential injections with Poly(I:C). Flow cytometry analysis of the BM 4 weeks post-Poly(I:C) revealed no significant difference in cell number or distribution of mature hematopoietic populations (myeloid, T-lymphoid, or B-lymphoid); however, the lineage negative cells, which include HPSCs, showed a decrease in total number upon loss of *Fbxo9* (Figure 3B-C, S3A-D). Further analysis of the HSPCs revealed that *Fbxo9* cKO results in increased stem and early progenitor cells (LSK), specifically the multipotent progenitors (Figure 3D-E, S3D). The cKit^+^ progenitor population (cKit^+^) decreased in total number due to a decrease in megakaryocyte-erythroid progenitors (Figure 3F). To determine whether loss of *Fbxo9* alters differentiation of HSPCs, we carried out a colony forming cell assay and found that LSKs derived from *Fbxo9^+/−^* and *Fbxo9^−/−^* gave rise to fewer colonies, showing decreased ability of HSPCs to differentiate and proliferate in response to cytokine stimulation (Figure 3G). We further analyzed these populations for cell cycle changes and found an increased number of cKit^+^ cells and LSKs in a quiescent G_0_ state (Figure 3H, S3D). Together these results suggest that HSPC populations are more quiescent and have decreased differentiation.

**Figure 3.**
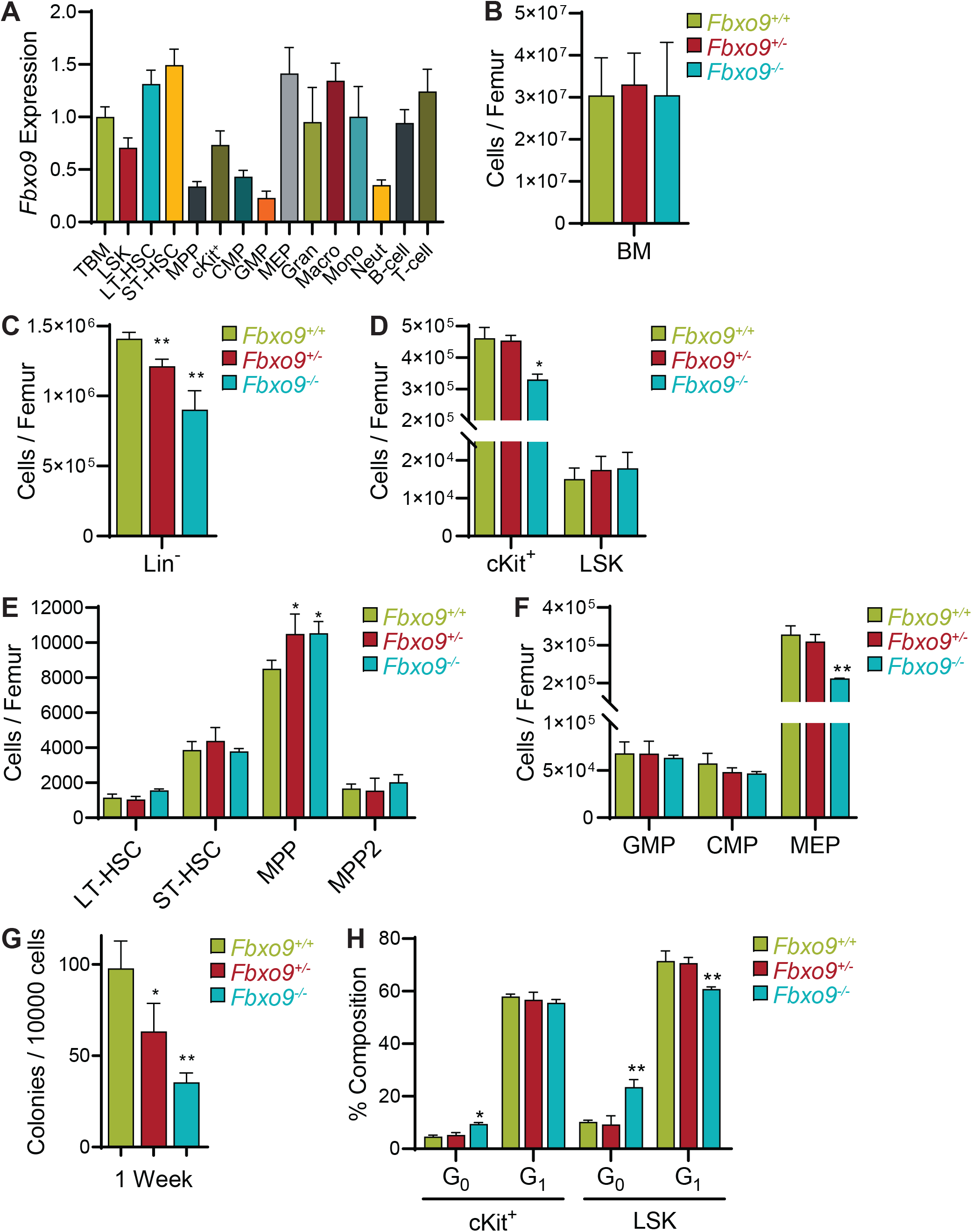
Loss of *Fbxo9* alters murine HSPC function. A Analysis of *Fbxo9* expression in WT murine hematopoietic lineages shows high expression in the long-term stem cells (LT-HSCs; Lin^−^cKit^+^Sca-1^+^CD150^+^CD48^−^), short-term stem cells (ST-HSCs; Lin^−^cKit^+^Sca-1^+^CD150^−^CD48^−^), megakaryocyte-erythroid progenitors (MEPs; Lin^−^cKit^+^Sca-1^−^CD34^−^CD16/32^−^), and macrophages (CD11b^+^Gr1^−^) when compared to total BM. B Bar graph of cell counts from BM extracted from the right femur of sacrificed mice. C-F Bar graphs showing the cell count of Lin^−^, cKit^+^ (Lin^−^ cKit^+^Sca-1^−^), LSKs (Lin^−^cKit^+^Sca-1^+^), LT-HSC, ST-HSC, multipotent progenitors (MPP; Lin^−^ cKit^+^Sca-1^+^CD150^−^CD48^+^), MPP2 (Lin^−^cKit^+^Sca-1^+^CD150^+^CD48^+^), granulocyte-macrophage progenitors (GMP; Lin^−^cKit^+^Sca-1^−^CD34+CD16/32^hi^), common myeloid progenitor (CMP; Lin^−^ cKit^+^Sca-1^−^CD34^+^CD16/32^lo^), and MEP compartments in the BM of mice of the indicated genotypes. G Bar graph of colonies per 10000 cells plated in methyl cellulose. H Bar graph showing the percentages of cells in G_0_ and G_1_ (for all data shown, bar graphs are mean ± standard deviation, n = 3, * p < 0.05, ** p < 0.01).

### Deletion of Fbxo9 leads to an aggressive and immature AML phenotype

The lowest expression of *FBXO9* was found in the inv(16) subtype of AML. Inv(16)/t(16;16) arises from an inversion or translocation within chromosome 16 that fuses the genes for core binding factor beta (*Cbfb*) and smooth muscle myosin heavy chain (*MYH11*)^31^. The resulting CBFB-SMMHC fusion protein alters Runt-related protein 1 (RUNX1, formerly AML1) activity, a transcription factor important in hematopoietic regulation, which leads to a block in differentiation of myeloid progenitor cells^32,33^. To study the role of *Fbxo9* in inv(16) AML, we crossed our *Mx^Cre^Fbxo9^f/f^* mouse with the floxed allele of *Cbfb-MYH11* (*Cbfb^+/56M^*) which allows for inducible expression of CFBF-SMMHC following administration of Poly(I:C)^32^. In mice, expression of the *Cbfb^+/56M^* allele is sufficient to initiate the formation of AML with a median survival of approximately 20 weeks^32^. We confirmed the presence of *Cbfb^56M^* within the offspring prior to Poly(I:C) treatment and confirmed deletion of exon 4 and expression of *Cbfb-MYH11* following Poly(I:C) by PCR (Figure 2B).

To monitor disease onset, we performed flow cytometry on peripheral blood (PB) beginning 3 weeks post-Poly(I:C) injection. Cell surface markers for mature myeloid cells Gr1 and CD11b were analyzed along with cKit to identify immature AML blasts. Analysis revealed that *Cbfb^+/56M^Fbxo9^+/+^* mice tend to have early (2-3 weeks) expression of cKit^+^ cells that diminishes at weeks 6 and 9. Although not significantly different, the kinetics of the cKit^+^ tumor population following *Fbxo9* deletion conversely expands at weeks 6 and 9 (Figure 4A). In addition, we found that most of the mice developed AML within the expected latency period, though the *Cbfb^+/56M^Fbxo9^+/−^* group had a significantly shorter time of survival with a median survival of 13 weeks compared to 17 weeks in *Cbfb^+/56M^Fbxo9^−/−^* and 20 weeks in *Cbfb^+/56M^Fbxo9^+/+^* cohorts (Figure 4B).

**Figure 4.**
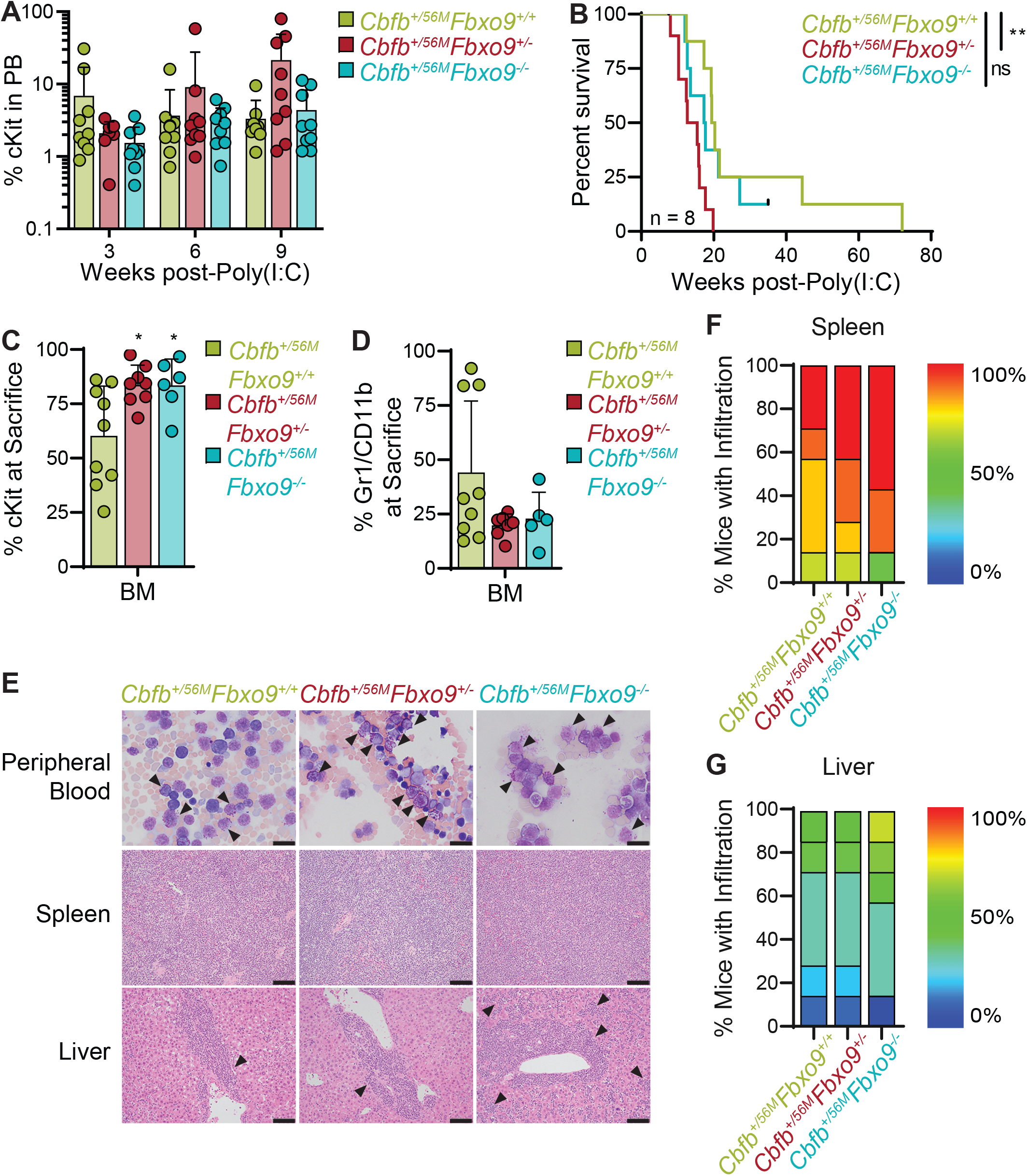
Loss of *Fbxo9* accelerates inv(16) AML and causes a more aggressive disease phenotype. A Analysis of the PB of mice following initiation of inv(16) AML shows that the cKit^+^ tumor population expands more rapidly in *Fbxo9* KO mice (mean ± standard deviation). B Kaplan-Meier survival curves of Poly(I:C) treated mice shows that loss of *Fbxo9* reduced time of survival (n = 8). C-D Bar graphs (mean ± standard deviation) representing the percentage of C cKit^+^ tumor cells or D Gr1^+^/CD11b^+^ cells in the BM of mice at time of sacrifice. E Representative images of Wright-Giemsa-stained peripheral blood and hematoxylin-eosin (H&E)-stained spleen and liver sections of mice with the indicated genotypes at time of sacrifice. Bars, 20 μm. F-G Quantification of the infiltration in the F spleen and G liver where red represents 100% infiltration and complete effacement, while blue represents no infiltration with normal tissue architecture (n = 7) (* p < 0.05, ** p < 0.01).

Consistent with previously reported data, ~80% of mice in the *Cbfb^+/56M^Fbxo9^+/+^* control group had tumors with a predominantly blast-like population, expressing the cell surface marker cKit, and ~20% expressed the more mature cell surface markers Gr1 and CD11b^32^. Upon deletion of *Fbxo9*, all of the mice from both *Cbfb^+/56M^Fbxo9^−/−^* and *Cbfb^+/56M^Fbxo9^+/−^* cohorts developed blast-like tumors expressing cKit on the cell surface and lacking Gr1/CD11b expression, suggesting a more immature phenotype (Figure 4C-D, S4A). Consequently, cKO mice had greater expression of cKit in BM upon sacrifice indicating a greater tumor burden in the BM upon loss of one or both *Fbxo9* alleles (Figure 4C). These data showed that *Cbfb^+/56M^Fbxo9^+/−^* results in a shorter time of survival and both *Cbfb^+/56M^Fbxo9^+/−^* and *Cbfb^+/56M^Fbxo9^−/−^* give rise to tumors with a more immature phenotype.

In addition, we analyzed secondary organs for signs of infiltration. The spleens from all groups had splenomegaly (Figure S4B). H&E staining of spleen infiltration demonstrated that ~60% of *Cbfb^+/56M^Fbxo9^−/−^* mice had complete effacement of the spleen architecture compared to only 30% of *Cbfb^+/56M^Fbxo9^+/+^* mice (Figure 4E-F). Likewise, in the liver *Cbfb^+/56M^Fbxo9^−/−^* mice had ~50% infiltration in half the cohort indicating a more aggressive disease even though they had a similar time of survival to *Cbfb^+/56M^Fbxo9^+/+^* mice (Figure 4G). These findings suggest that loss of *Fbxo9* leads to increased infiltration of spleen and liver.

### Transplanted tumors with reduced Fbxo9 lead to rapid onset of disease

Development of primary AML tumors demonstrated that loss of *Fbxo9* alters the AML phenotype. To determine whether *Fbxo9* is acting in a tumor initiating or tumor-promoting fashion, we carried out a secondary transplantation. Tumor cells from spleens were injected into sub-lethally irradiated recipient mice. To eliminate bias from tumor phenotype, we selected primary splenic tumor cells with a cKit^+^ phenotype (Figure 5A). All tumors selected contained >90% tumor burden with the exception of one *Cbfb^+/56M^Fbxo9^+/+^* tumor with only 75% cKit^+^ spleen cells. The mouse was selected due to aggressive nature of the tumor which resulted in the shortest survival from the *Cbfb^+/56M^Fbxo9^+/+^* cohort.

**Figure 5.**
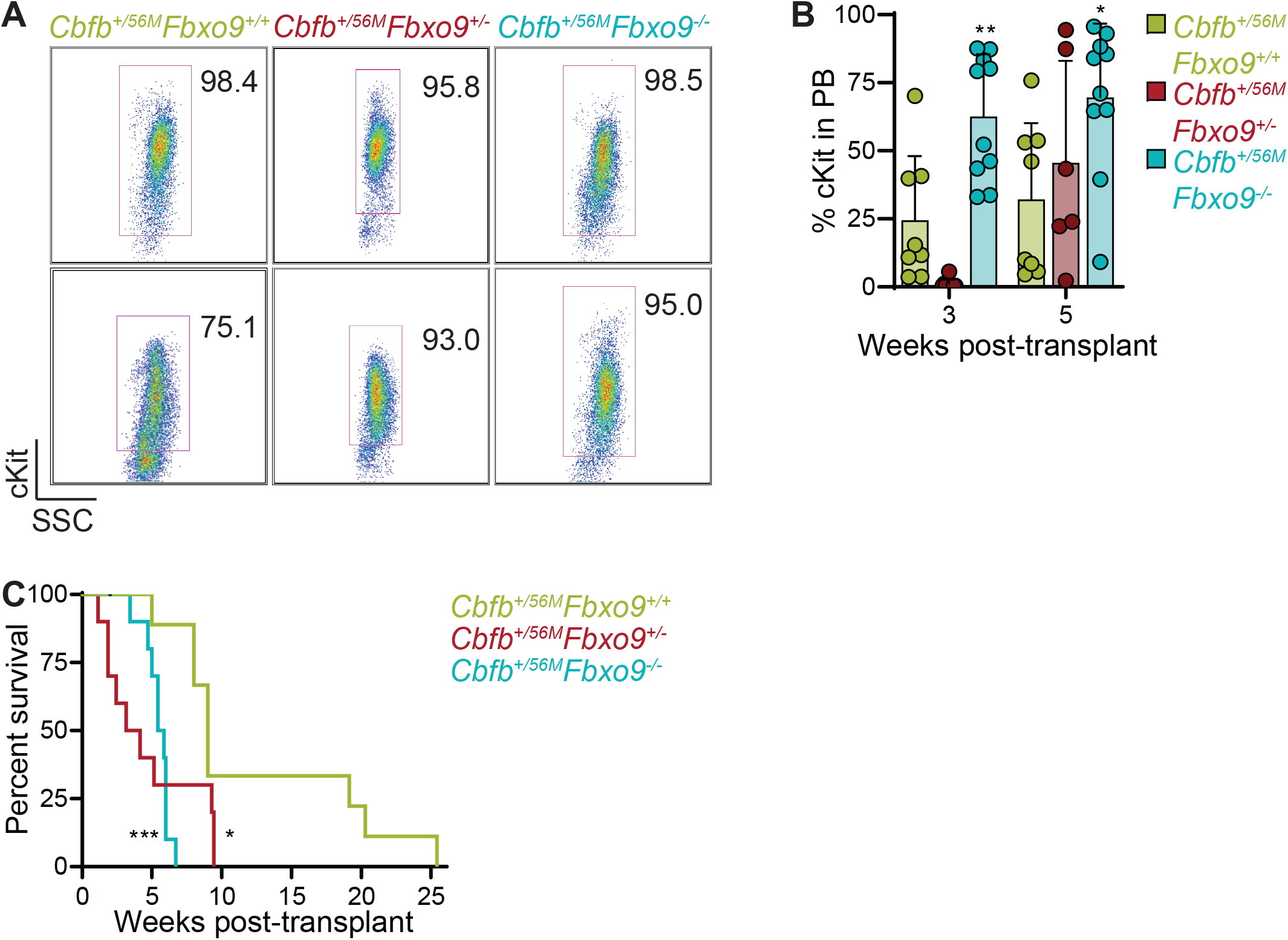
Transplant of inv(16) spleen tumor cells with lower expression of *Fbxo9* develop more aggressively. A FACS plots of spleen tumor cells transplanted into sub-lethally irradiated host mice. B Bar graph (mean ± standard deviation) representing the percentage of cKit+ cells in the PB of transplant mice. C Overall survival of mice transplanted with tumors of the indicated genotypes (n = 10, * p < 0.05, ** p < 0.01, *** p < 0.001).

We followed the development of AML and by 5 weeks post-transplant *Cbfb^+/56M^Fbxo9^−/−^* mice had on average 69.6% tumor PB indicating that tumors with decreased *Fbxo9* develop more rapidly (Figure 5B, S5A). The rapid expansion of the tumor population led to decreased time of survival in *Cbfb^+/56M^Fbxo9^+/−^* and *Cbfb^+/56M^Fbxo9^−/−^* cohorts compared to *Cbfb^+/56M^Fbxo9^+/+^* mice (Figure 5C). Upon sacrifice, analysis of the BM, spleen, and PB by flow cytometry showed no difference in tumor burden between the three cohorts (Figure S5B). Secondary transplantation confirms that tumors lacking *Fbxo9* are aggressive and lead to rapid progression of AML.

### Loss of Fbxo9 leads to up-regulation of proteins associated with tumorigenicity

To identify potential *Fbxo9* substrates and protein alterations in AML, we performed quantitative MS on splenic tumor cells. We labeled proteins isolated with tandem mass tags, combined equal concentrations from each sample, and analyzed by MS (LC-MS/MS) (Figure 6A). Mass spectrometry identified 18696 unique peptides representing 3580 proteins, 3247 of which were quantifiable (Figure 6B). To quantify protein changes, we compared only proteins with ≥3 unique peptides. Between *Cbfb^+/56M^Fbxo9^+/−^* and *Cbfb^+/56M^Fbxo9^−/−^* cohorts, 118 proteins were significantly up-regulated (p<0.05, fold change ≥1.3) and 86 proteins were significantly down-regulated (p<0.05, fold change ≤0.7) in one or both cohorts (Figure 6C, Table S1).

**Figure 6.**
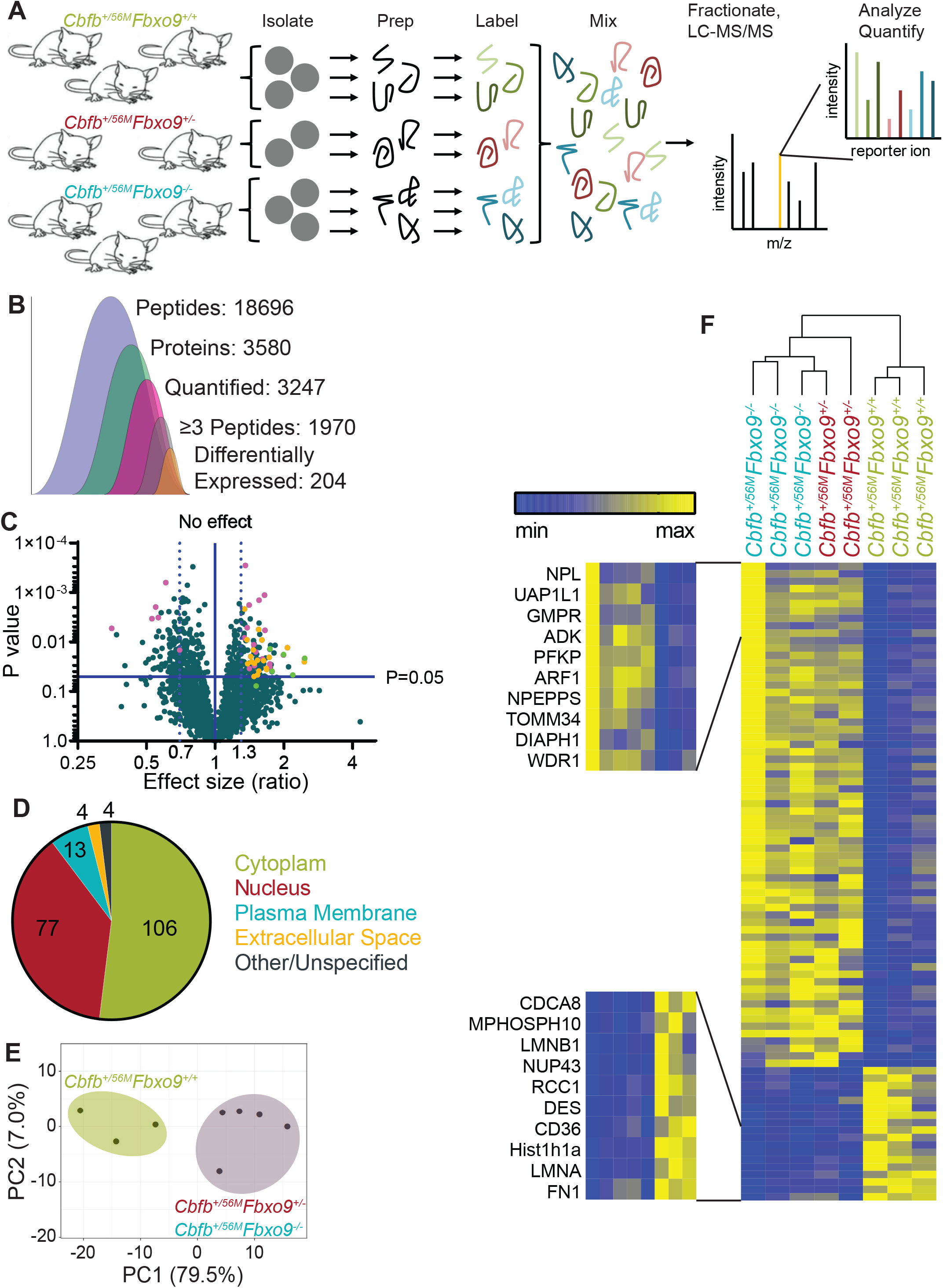
Tumors lacking *Fbxo9* are more similar than tumors with WT expression. A Schematic of preparation for TMT MS using tumors isolated from *Cbfb^+/56M^Fbxo9^+/+^*, *Cbfb^+/56M^Fbxo9^+/−^*, and *Cbfb^+/56M^Fbxo9^−/−^* mice (n = 3). B Quantification of the identified peptides and proteins. C Volcano plot of the fold change *Cbfb^+/56M^Fbxo9^−/−^/Cbfb^+/56M^Fbxo9^+/+^* samples for significantly up-regulated and significantly down-regulated proteins (pink, significantly upregulated in *Cbfb^+/56M^Fbxo9^+/−^/Cbfb^+/56M^Fbxo9^+/+^*; yellow, proteins participating in proteasome-dependent pathway; green, top up-regulated proteins associated with tumorigenicity). D Pie chart of localizations for the differentially expressed proteins identified in either the *Cbfb^+/56M^Fbxo9^+/−^*, *Cbfb^+/56M^Fbxo9^−/−^*, or both cohorts. E PCA plot using components 1 and 2 showing clustering of *Cbfb^+/56M^Fbxo9^+/+^* tumors compared to *Cbfb^+/56M^Fbxo9^+/−^* and *Cbfb^+/56M^Fbxo9^−/−^* tumors. F Heatmap with hierarchical clustering of the significantly up-regulated (p < 0.05, ≥ 1.3 fold increase over WT) and down-regulated (p < 0.05, ≤ 0.7 fold decrease from WT).

The majority of these differentially expressed proteins localized to the cytoplasm where FBXO9 is expressed (Figure 6D). To determine whether tumors with different genotypes show distinct patterns of expression, we carried out a principle component analysis and hierarchical clustering and found that 3 independent tumors with WT expression of *Fbxo9* clustered together while heterozygous and homozygous cKO tumors clustered together, indicating they are more similar to each other than to *Cbfb^+/56M^Fbxo9^+/+^* tumors (Figure 6E-F). Interestingly, a number of the top up-regulated proteins identified (PFKP, ADK, ARF1, TOMM34, DIAPH1, and WDR1) have been shown to participate in cancer by increasing cell growth and metastasis or are biomarkers for poor outcome (Figure 6F)^34–39^.

### Loss of Fbxo9 correlates with increased proteasome activity

The proteins overexpressed upon loss of *Fbxo9* were also enriched for proteins associated with proteasome-mediated pathways such as proteolysis and ubiquitin- or proteasome-dependent catabolism (Figure 7A). Increased proteasome activity has previously been implicated in cancer aggression and proteasome inhibitors have been approved as a treatment option^40^. Considering this strong correlation between cancer and proteasome activity, we confirmed proteasome component overexpression by western blot and observed an increase in differentially expressed proteasome components in tumors lacking *Fbxo9* (Figure 7B-C). To confirm that proteasome component expression correlates with increased activity *in vitro*, we performed a proteasome activity assay comparing our tumor groups. This confirmed that not only does loss of *Fbxo9* result in increased proteasome component expression, but that this expression correlates with increased proteasome activity in the *Cbfb^+/56M^Fbxo9^+/−^* and *Cbfb^+/56M^Fbxo9^−/−^* tumors (Figure 7D). To further elucidate the effect of loss of *Fbxo9* in AML, we treated cultured tumor cells with varying concentrations of proteasome inhibitor bortezomib (Figure 7E). Bortezomib treatment confirmed that *Cbfb^+/56M^Fbxo9^−/−^* tumor cells are more sensitive to proteasome inhibition than *Cbfb^+/56M^Fbxo9^+/+^* tumor cells with IC_50_ calculations of 10.03nM and 11.76nM, respectively. Furthermore, analysis of apoptosis and cell death demonstrated that *Cbfb^+/56M^Fbxo9^−/−^* tumor cells are more sensitive to treatment with bortezomib than *Cbfb^+/56M^Fbxo9^+/+^* tumor cells (Figure 7F). These studies provide evidence that loss of *Fbxo9* leads to increased proteasome activity and sensitivity to proteasome inhibitors like bortezomib.

**Figure 7.**
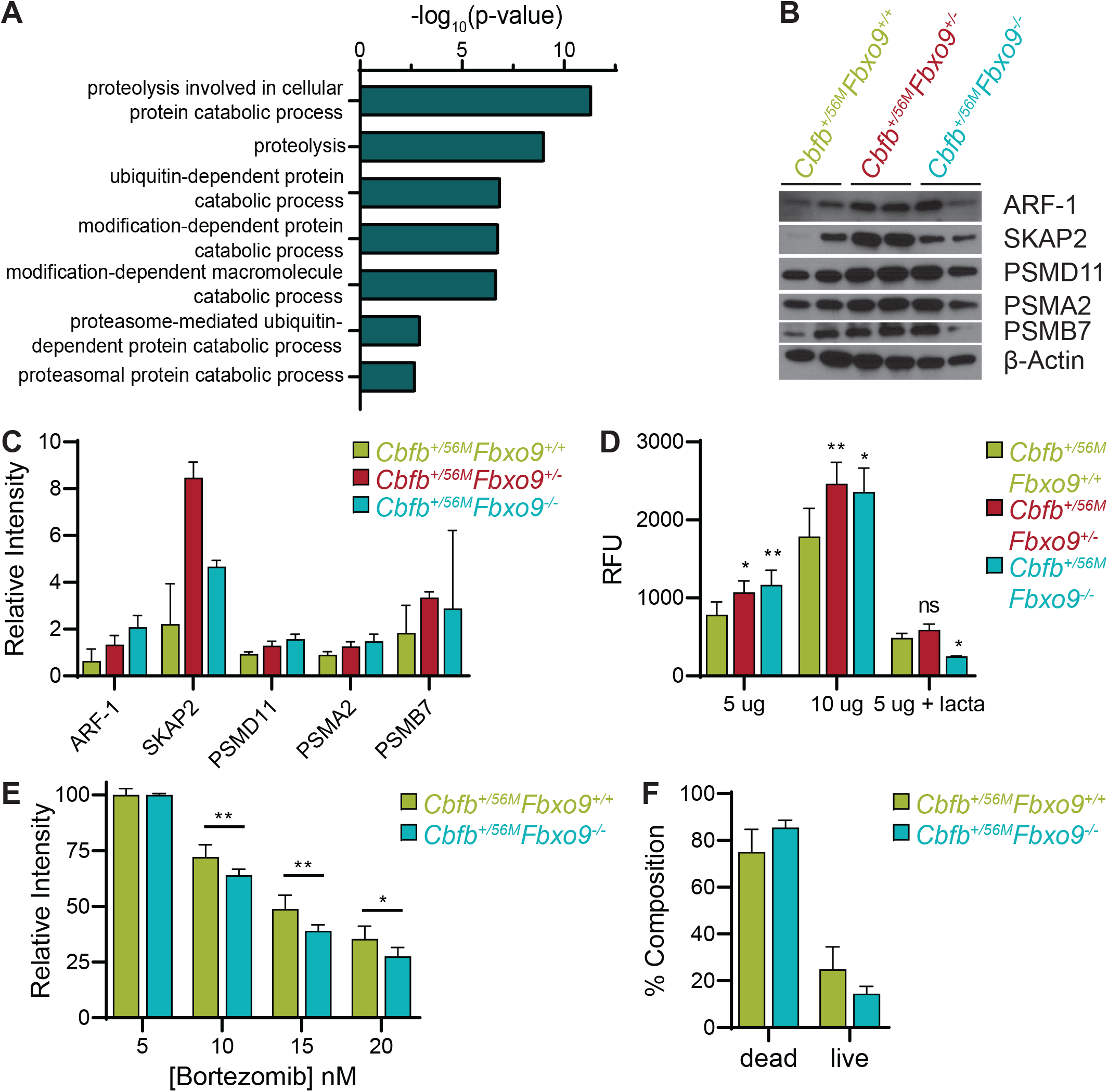
Knockout of *Fbxo9* results in an increase in proteasome activity and differing response to proteasome inhibition *in vitro*. A Gene ontology analysis showing pathways known to be associated with significantly up-regulated (p < 0.05, ≥ 1.3 fold change) proteins using DAVID 6.8 software. For up-regulated proteins, pathways included have ≥ 10 associated proteins and p < 0.01. B Western Blot analysis of identified up-regulated proteins and proteasome components for expression in murine inv(16) AML tumors with the indicated genotypes. C Bar graph (mean ± standard deviation) quantifying relative expression of the proteins compared to β-actin expression. D Bar graph (mean ± standard deviation) of proteasome activity in cultured tumor cells using the indicated amounts of protein, proteasome activity confirmed by treatment with the proteasome inhibitor lactacystin. E Bar graph (mean ± standard deviation) showing survival of *Cbfb^+/56M^Fbxo9^+/+^* and *Cbfb^+/56M^Fbxo9^−/−^* tumor cells after 24 h culture with increasing doses of bortezomib. F Bar graph (mean ± standard deviation) quantifying cell death of tumors cultured 16 h with 20 nM bortezomib, cell death analyzed by flow cytometry using Annexin-V and 7AAD (for all data shown n = 3, * p < 0.05, ** p < 0.01).

## DISCUSSION

The molecular pathogenesis of AML has yet to be fully defined, though many acquired molecular and cytogenetic abnormalities have been identified that lead to leukemogenesis^41^. The majority of these alterations are thought to occur in the HSPCs and disease progression is often thought to progress through the dysregulation of normal cellular mechanisms^42^. As FBXO9 is an E3 ligase highly expressed in HSPCs (the tumor initiating population) it is imperative to understand the resulting changes in HSPC function upon loss of its expression. We first report that loss of *Fbxo9* correlates with a decrease in the Lin^−^ cells of the BM but an increase in the LSKs within that compartment. Second, we show that the increase is due to an expansion of quiescent cells in G_0_ phase of the cell cycle which leads to a decrease in colony formation when these HSPCs are in culture. To further understand the mechanistic changes associated with disease initiation and progression, it is essential to determine the role FBXO9 plays in the tumor initiating population.

Aberrant FBXO9 expression has previously been linked to disease progression in MM by tagging mTORC1 components for degradation^25^. In this context, its overexpression was shown to promote disease progression and FBXO9 has previously been classified as an oncogene. Contrary to its role in MM, our studies demonstrate that *FBXO9* expression is consistently decreased across AML subtypes and that reduction of its E3 ligase activity promotes the progression of AML and correlates with a shorter time of survival. Additionally, MS experiments did not show accumulation of the known substrates TEL2 and TTI1 identified in MM. This finding demonstrates that FBXO9 plays a context-specific role in cancer, acting as an oncogene in MM and a tumor suppressor in AML. While other F-box proteins have been shown to have solely oncogenic or tumor suppressor roles, FBXO9 is among a few select members of the family that can function in both capacities^16^. Furthermore, we found that loss of a single copy of *Fbxo9* was sufficient to cause increased aggressiveness in AML tumor cells. Considering that FBXO9 is an E3 ligase, we must elucidate the alterations that loss of this protein causes in the proteomic landscape of AML cells.

Through interrogation of the proteomic changes occurring in AML tumor cells upon loss of *Fbxo9*, we identified various proteasome components and proteasome-related proteins that were upregulated. Increased proteasome activity has been associated with aggressiveness in many cancer subtypes through *in vitro* and *in vivo* studies^43–46^. Ma *et al.* reported that AML patients have higher levels of 20S proteasome components compared to healthy controls but that this increase does not lead to increased chymotrypsin-like activity due to a reduction in expression of the 19S regulatory component^47^. Similarly, our data showed increased proteasome component expression of the 20S catalytic subunit (PSMA and PSMB) in *Cbfb^+/56M^Fbxo9^−/−^* compared to *Cbfb^+/56M^Fbxo9^+/+^*. Unlike the results produced by Ma *et al.*, we saw increased expression in the 19S (PSMD) components indicating that loss of *Fbxo9* could correspond with altered proteasome activity. Further analysis revealed that increased component expression did, indeed, lead to increased proteasome activity, suggesting a causal relationship.

Proteasome inhibition with drugs such as bortezomib has become standard of care for patients with MM and has been effectively utilized as a second-line therapy in treating mantle cell lymphoma and follicular lymphoma^48–50^. Clinical trials using bortezomib, either alone or in combination with other chemotherapies, have reported varying complete remission rates ranging from 0-80%^51–57^. To date, no correlation has been made between AML classification and patients who respond well to proteasome inhibition. Indeed, responses seem to be independent of AML subtype. Additionally, one of the main impediments to achieving complete remission in AML stems from the inability of current therapies to target the leukemic stem cells. Proteasome inhibitors have previously been effective in targeting this tumor-initiating population and the findings presented herein could lead to more efficacious clinical use of agents like bortezomib, particularly in targeting the leukemic stem cells^58,59^. Overall, we have identified *Fbxo9* as a tumor suppressor of AML and shown that loss of its expression leads to increased proteasome activity and sensitivity to proteasome inhibition, thus implying that *FBXO9* expression could be used as an indicator for patients who would respond well to proteasome inhibition.

## METHODS

### Transgenic mouse models

*Fbxo9* cKO mice were developed using Easi-CRISPR as previously published^29^ and bred with *Cbfb^+/56M^* [^29,32^] or Mx^cre^ mice purchased from Jackson Laboratories (#003556, Bar Harbor, ME, USA). PCR confirmed genotype and expression (primers in supplemental materials and methods). To induce cKO of *Fbxo9* and expression of *Cbfb-MYH11*, 6-8 week-old floxed mice and littermate controls received three intraperitoneal Poly(I:C) injections every other day at a dose of 10μg per gram body weight (Invivogen). Procedures performed were approved by the Institutional Animal Care and Use Committee of the University of Nebraska Medical Center in accordance with NIH guidelines.

### Flow Cytometry analysis

For flow cytometry analyses, PB was extracted from the tail vein and RBCs were lysed with 500μL ACK lysing buffer. Upon sacrifice, spleen and BM cells were strained through 0.45μm strainer and treated with ACK lysing buffer. Spleen, BM, and PB cells were stained for 1h on ice in the dark in 3% FBS in PBS (antibodies in supplemental materials and methods). For cell cycle analysis, cells were fixed and permeabilized following Biolegend intracellular staining protocol and stained with Ki67 and DAPI using 1μL DAPI per sample.

### Colony Forming Cell (CFC) Assay

Fresh or culture progeny from total BM were counted and 5000 cells/well of a 24-well plate were resuspended in Methocult (M3434, Stem Cell Technologies, Vancouver, BC, Canada). Colonies were counted at day 7, resuspended, and replated as before.

### Histological Staining

Mouse organs were fixed in 10% (vol/vol) buffered formalin phosphate for 24h and stored in 70% ethanol. Sections were stained with H&E using standard protocols. The slides were evaluated and graded by treatment group by a pathologist for leukemia infiltration.

### Western Blot Analysis

For western blot analysis, samples were lysed in an IP lysis buffer (20mM Tris pH7.5, 150mM NaCl, 1mM EDTA) containing 1X Halt Protease and Phosphatase Inhibitor Cocktail (ThermoFisher, Waltham, MA, USA) and 10mM NEM. Membranes were blocked in 5% milk. Antibodies were prepared in 5% BSA as indicated in supplemental materials and methods.

Horse Radish Peroxidase conjugated secondary antibodies (Jackson Laboratory) were prepared in 5% milk as indicated in supplemental materials and methods.

### RNA extraction and quantitative RT-PCR

Total RNA was harvested from BM cells using the RNeasy Kit (QIAGEN, Hilden, Germany). Following extraction, total RNA was used for cDNA synthesis using the High Capacity RNA-to-cDNA Kit (ThermoFisher). cDNA was quantified by measuring absorbance at A280nm and qRT-PCR was carried out on equal concentrations of cDNA from each sample using the iTaq Universal SYBR Green Supermix (BioRad, Hercules, CA, USA). Primers in supplemental materials and methods.

### TMT labeling and Mass Spectrometry

For global proteome quantification, splenic tumor cells were isolated as described above from 2-3 mice per genotype. Samples were prepared and TMT-labeled per manufacturer’s protocol (ThermoFisher TMT10plex Mass Tag Labeling Kits). Following TMT labeling, acetonitrile was removed by speedvac and samples were resuspended in 0.1% trifluoroacetic acid. Sample cleanup with C18 tips was performed per manufacturer’s protocol (Pierce). Sample concentrations were re-quantified (Pierce Quantitative Colorimetric Peptide Assay kit) and combined in equal concentration. Following combination, samples were dried by speedvac and fractionated by ThermoFisher high pH reverse phase fractionation kit following manufacturer’s protocol for TMT. Resulting fractions were dried in a speedvac and resuspended in 0.1% Formic Acid for MS analysis (see supplemental methods). Data are available via ProteomeXchange with identifier PXD014387^60^.

### Proteasome Activity Assay

Tumor cells from isolated spleen tissue (2×10^6^) were lysed in 0.5% NP-40 in PBS for 10min on ice with vortexing. 5-10 μg protein lysates were cultured and proteasome activity measured using the 20S Proteasome Activity Assay kit (APT280, Millipore, Billerica, MA, USA) per manufacturer’s protocol.

### MTT Assay

2×10^5^ cells were plated into each well of a 96-well plate and cultured for 24h in 100μL tumor growth medium (StemSpan SFEM, 5% Penn/Strep, 5% Lipid mixture, 5% Glutamate, 20ng/mL SCF, 10ng/mL IL-6, 10ng/mL IL-3) containing DMSO or varying concentrations of bortezomib. Following culture, CellTiter 96 AQ_ueous_ One Solution Cell Proliferation Assay (MTS) (Promega, Madison, WI, USA) was performed per manufacturer’s protocol.

### Statistical analysis

All experiments were performed in triplicate unless noted and statistical analyses were performed using unpaired two-tailed Student’s t-test assuming experimental samples of equal variance. * p-value<0.05, ** p-value<0.01, *** p-value<0.001, **** p-value<0.0001.

## AUTHOR CONTRIBUTIONS

R.W.H., and S.M.B. conceived and designed the experiments. R.W.H., K.J.W., M.C., S.A.S., H.V., and S.M.B preformed experiments and analysis. R.W.H., and S.M.B. wrote the manuscript. R.K.H. provided technical and material support. C.A. provided hematopathology expertise. All authors reviewed the manuscript before submission.

## ACKNOWLEDGEMENTS

We would like to thank the UNMC Flow Cytometry Research Facility, UNMC Mouse Genome Engineering Core Facility, and UNMC Mass Spectrometry and Proteomics Core Facility for expert assistance. The core facilities are administrated through the Office of the Vice Chancellor for Research and supported by state funds from the Nebraska Research Initiative (NRI) and The Fred and Pamela Buffett Cancer Center’s National Cancer Institute Cancer Support Grant. S.M.B. is supported by the National Institutes of Health (P20GM121316), and ACS institutional grant. R.K.H., and S.M.B are supported by the UNMC Pediatric Cancer Group and by National Institute of General Medical Sciences of the National Institutes of Health under grant number p30GM106397. R.W.H. is supported by the UNMC NIH training grant (5T32CA009476-23). This publication was supported by the Fred & Pamela Buffett Cancer Center Support Grant from the National Cancer Institute under award number P30 CA036727.

