## Supplemental Material for "Loss of FBXO9 enhances proteasome activity and promotes aggressiveness in acute myeloid leukemia"

**SUPPLEMENTAL MATERIALS AND METHODS**

| Genotyping Primer | Sequence |
|---|---|
| Fbxo9 5' F | 5' – GTC TCT TCG AGG CAG AAC GTA TC – 3' |
| Fbxo9 5' R | 5' – GCA ACG GGA TCA GAT GTC CAA TG – 3' |
| Fbxo9 3' F | 5' – GCC TGA GAC TGA AAG ATC CTT GC – 3' |
| Fbxo9 3' R | 5' – CAG GCT TAC AAA GAG GGC CTA CG – 3' |
| Hygro F | 5' – CCA TCG TCG AGA TCC AGA CAT – 3' |
| Hygro R | 5' – GTA TAT GCT CCG CAT TGG TCT TG – 3' |
| Generic Cre F | 5' – GCG GTC TGG CAG TAA AAA CTA TC – 3' |
| Generic Cre R | 5' – GTG AAA CAG CAT TGC TGT CAC TT – 3' |
| Inv Post F | 5' – CCC CCG TAC TGT GTG TGT CT – 3' |
| Inv Post R | 5' – GCC AGA CGG GTC AAC AAT AC – 3' |

| qRT-PCR Primer | Sequence |
|---|---|
| Gapdh F | 5' – CAT GGC CTT CCG TGT TCC TA – 3' |
| Gapdh R | 5' – CTG GTC CTC AGT GTA GCC CAA – 3' |
| Fbxo9 Exon 4 F | 5' – AGA AGC TCT ATG CTG AAA GCA G – 3' |
| Fbxo9 Exon 4 R | 5' – CAT CGC CCT ACG GTA GAA CT – 3' |
| Fbxo9 Exon 2-3 F | 5' – ATG AGA GTC CGG CTG AGA GA – 3' |
| Fbxo9 Exon 2-3 R | 5' – AGA GCT TCT TCC TGC TCT GC – 3' |

**Western Blot Antibodies:**

| Antibody | Company | Ref# |
|---|---|---|
| ARF-1 | Proteintech | 10790-1-AP |
| B-actin-HRP | Santa Cruz | sc-47778 |
| FLAG-HRP | Proteintech | 12926-1-AP |
| PSMA2 | Bethyl | A303-816A |
| PSMB7 | Bethyl | A303-848A |
| PSMD11 | Cell Signaling | 14303S |

**Flow Cytometry Antibodies**

| Antibody | Fluorochrome | Clone | Company |
|---|---|---|---|
| 7-AAD | n/a | n/a | eBioscience |
| Annexin V | PE | n/a | BioLegend |
| B220 | PacBlue, BV510, APCcy7, APC, Biotin, PE | RA3-6B2 | BioLegend |
| CD4 | APCcy7, APC, Biotin, PE | CK1.5 | BioLegend |
| CD8 | APCcy7, APC, Biotin, PE | 53-6.7 | BioLegend, eBioscience |
| CD11b | FITC, APCcy7, APC, Biotin, PE | M1/70 | BioLegend |
| CD16/32 | FITC, APC | 93 | BioLegend |
| CD34 | APC, PE | HM34 | BioLegend |
| CD48 | FITC | HM48-1 | BioLegend |
| CD131 | PE | n/a | BD Pharmingen |
| CD150 | BV510, BV785 | TC15-12F12.2 | BioLegend |
| cKit | APC, BV421, PacBlue | 2B8 | BioLegend |
| Gr-1 | APCcy7, APC, Biotin, PE, BV421 | RB6-8C5 | BioLegend |
| Ki67 | FITC | 16A8 | BioLegend |
| Sca-1 | PE, PEcy7 | D7 | BioLegend, eBioscience |
| Strep | FITC | n/a | BioLegend |
| Ter-119 | APCcy7, PacBlue, APC, Biotin, PE | TER-119 | BioLegend |

**Mass Spectrometry Method:**

Samples were loaded onto trap column Acclaim PepMap 100 75 µm x 2 cm C18 LC Columns (Thermo Scientific™) at flow rate of 5 µl/min then separated with a Thermo RSLC Ultimate 3000 (Thermo Scientific™) from 5-20% solvent B (0.1% FA in 80% ACN) from 10-98 minutes at 300 nL/min and 50 ºC with a 120 minutes total run time for fractions one and two. For fractions three to six, solvent B was used at 5-45% for the same duration. Eluted peptides were analyzed by a Thermo Orbitrap Fusion Lumos Tribrid (Thermo Scientific™) mass spectrometer in a data dependent acquisition mode using synchronous precursor selection method. A survey full scan MS (from m/z 375-1500) was acquired in the Orbitrap with a resolution of 120000. The AGC target for MS2 in iontrap was set as $1 \times 10^4$ and ion filling time set as 150ms and fragmented using CID fragmentation with 35% normalized collision energy. The AGC target for MS3 in orbitrap was set as $1 \times 10^5$ and ion filling time set as 200 ms with a scan range of 100-500 and fragmented using HCD with 65% normalized collision energy. Protein identification was performed using proteome discoverer software version 2.2 (Thermo Fisher Scientific) by searching MS/MS data against the UniProt mouse protein database. The search was set up for full tryptic peptides with a maximum of 2 missed cleavage sites. Oxidation, TMT6plex of the amino terminus, GG and GGQ ubiquitination, phosphorylation, and acetylation were included as variable modifications and carbamidomethylation and TMT6plex of the amino terminus were set as fixed modifications. The precursor mass tolerance threshold was set at 10 ppm for a maximum fragment mass error of 0.6 Da with a minimum peptide length of 6 and a maximum peptide length of 144. The significance threshold of the ion score was calculated based on a false discovery rate calculated using the percolator node. Protein accessions were put into Ingenuity Pathway Analysis (QIAGEN Inc.) to identify gene symbols and localizations. Gene ontology pathway analysis was performed using DAVID Bioinformatics Database 6.8 using the functional annotation tool.

**SUPPLEMENTARY FIGURES**

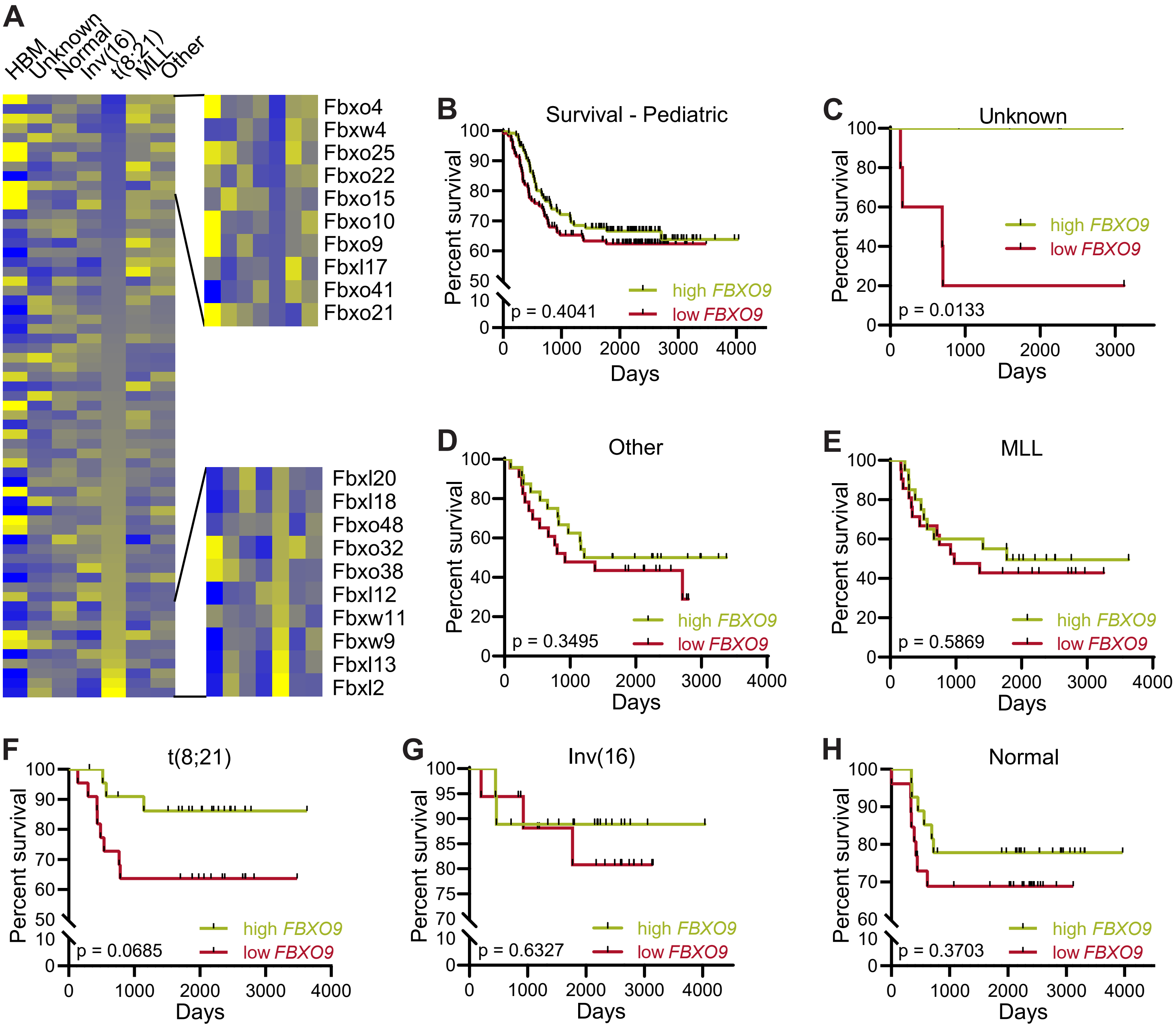


**Supplementary Figure S1 Pediatric patients with low *FBXO9* expression tend to have a shorter time of survival.** A Relative expression of F-box proteins in pediatric AML patients compared to healthy bone marrow (HBM). B-H Survival of pediatric patients of various subtypes grouped by *FBXO9* expression above (high) or below (low) the median.

**A** FBXO9 Translation Exon 3 Exon 4 Exon 5

Normal . . . QELAKEEKAREL . . . GALYEAIKFYRRA . . .

Mutant . . . QELAKEEK______________PSSSTVGR . . .

**B**

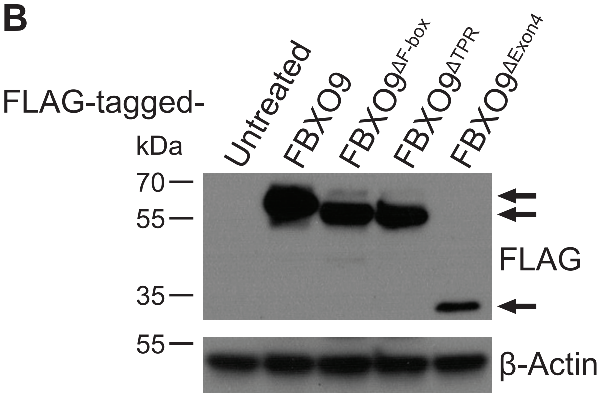


**Supplementary Figure S2 Deletion of *Fbxo9* exon 4 results in a frame shift and premature stop.** A Amino acid translation of sequenced cDNA from *Fbxo9*$^{+/+}$ and *Fbxo9*$^{-/-}$ mice. B Western blot expression of overexpressed flag-tagged FBXO9 with various deletion mutations.

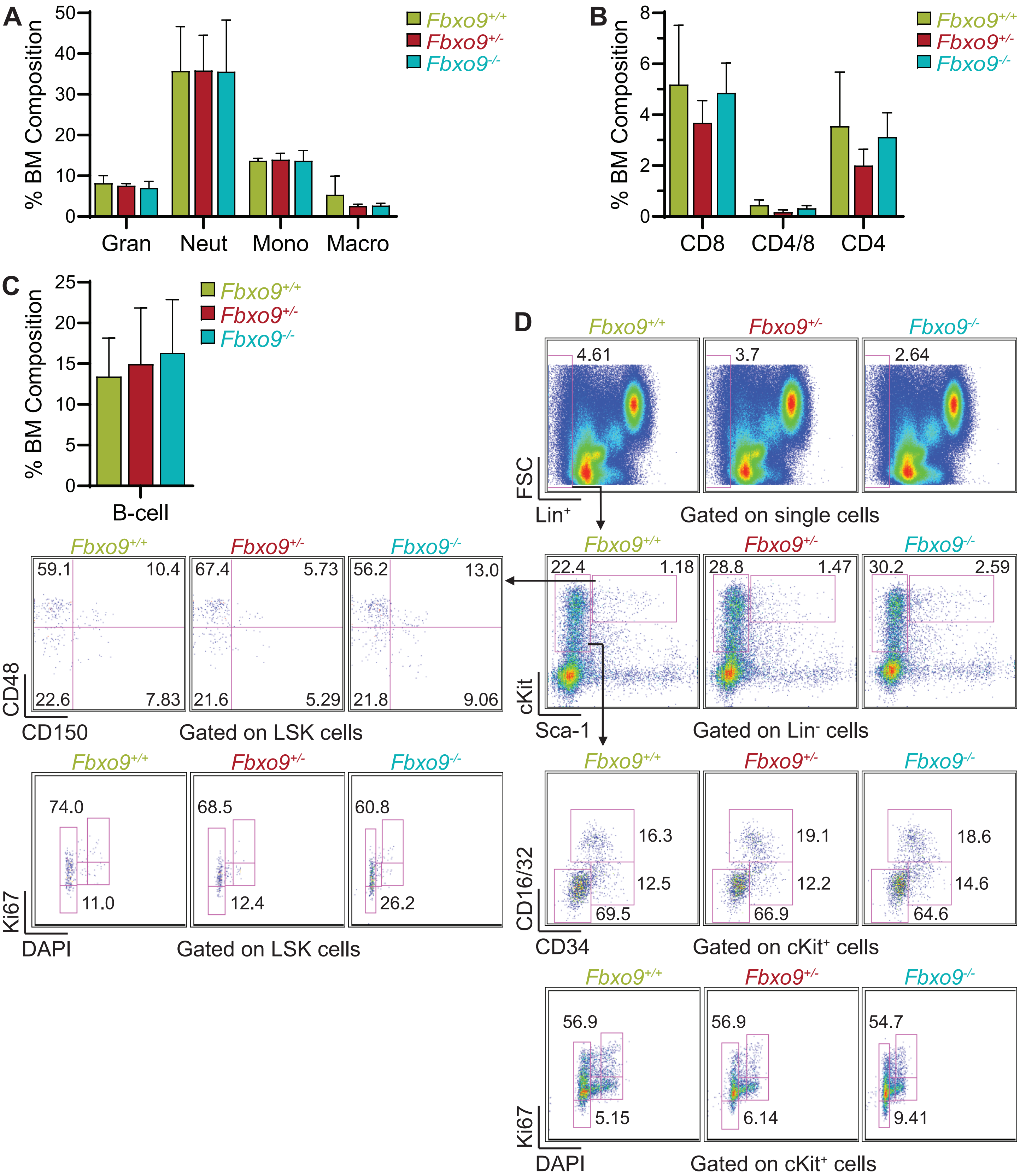


**Supplementary Figure S3 Loss of *Fbxo9* does not affect mature hematopoietic cell populations.** A-C Bar graph (mean ± standard deviation) of BM cell percentages of A myeloid cells, B T-cells, and C B-cells following treatment with Poly(I:C). D Flow cytometry gating strategy for isolating LSK and cKit$^{+}$ populations from total BM and analyzing for cell cycle stage.

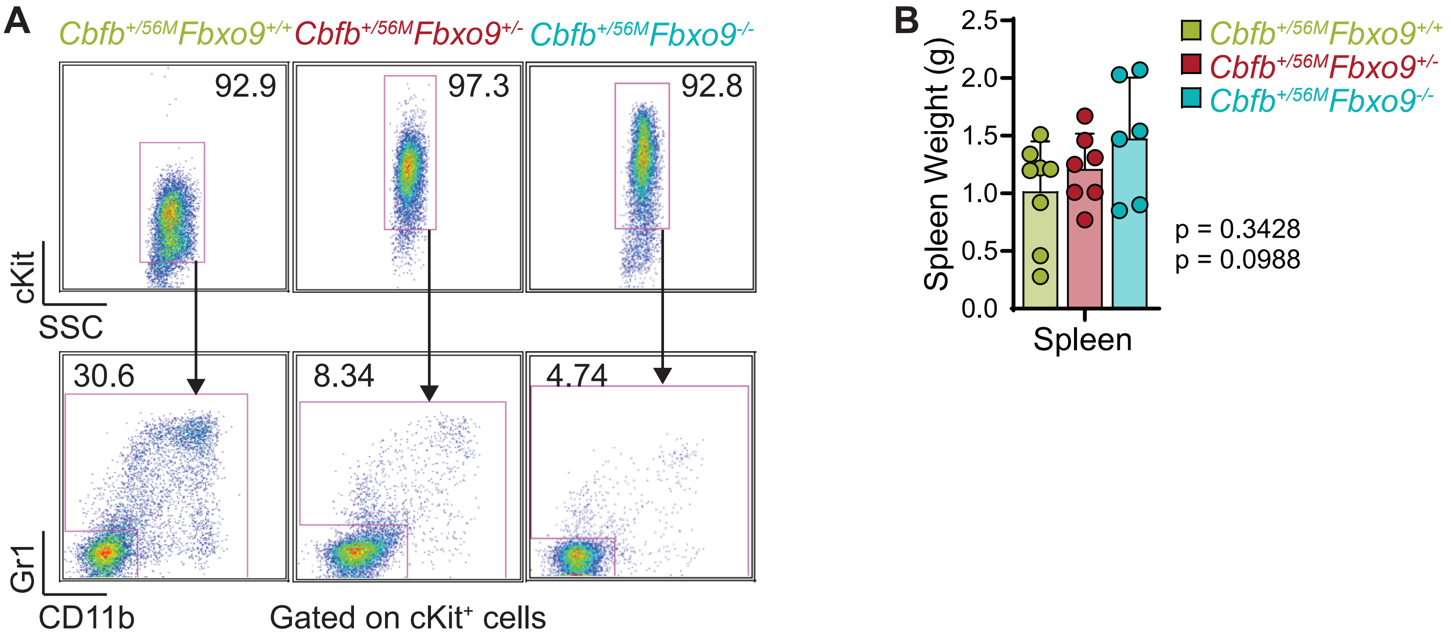


**Supplementary Figure S4 Mice with Inv(16) AML and *Fbxo9* cKO have a more aggressive and homogenous tumor.** A Representative FACS plots of tumor cells isolated from the BM of mice at time of sacrifice. Cells are first gated on total live singlets and subsequently on cKit⁺ cells which are further analyzed for expression of Gr1/CD11b. B Bar graph (mean ± standard deviation) of spleen weight of mice at time of sacrifice.

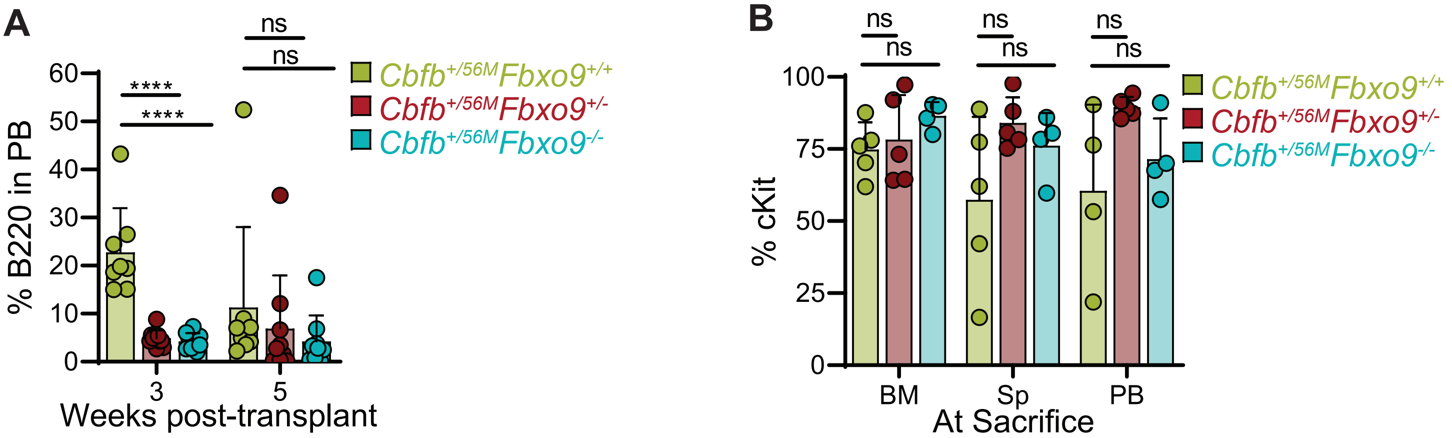


**Supplementary Figure S5 Mice with transplanted Inv(16) tumor cells show a more aggressive phenotype for cells lacking *Fbxo9* expression.** A Bar graph of B-cell (B220) expression in the PB following transplantation (mean ± standard deviation). B Bar graph of percentage of $cKit^{+}$ tumor cells in the BM, spleen (Sp), and PB at time of sacrifice (mean ± standard deviation, **** $p<0.0001$).
